# Enrichment and application of bacterial sialic acids containing polymers from the extracellular polymeric substances of “*Candidatus* Accumulibacter”

**DOI:** 10.1101/2022.09.16.508216

**Authors:** Sergio Tomás-Martínez, Le Min Chen, Martin Pabst, David G. Weissbrodt, Mark C.M. van Loosdrecht, Yuemei Lin

**Author notes:** Corresponding Author: Sergio Tomás-Martínez, Address: Department of Biotechnology, Delft University of Technology. Van der Maasweg, 9,2629 HZ, Delft, The Netherlands.

## Abstract

Pseudaminic and legionaminic acids are a subgroup of nonulosonic acids (NulOs) unique to bacterial species. There is a lack of advances in the study of these NulOs due to their complex synthesis and production. Recently, it was seen that “*Candidatus* Accumulibacter” can produce Pse or Leg analogues as part of its extracellular polymeric substances (EPS). In order to employ a “*Ca*. Accumulibacter” enrichment as production platform for bacterial sialic acids, it is necessary to determine which fractions of the EPS of “*Ca*. Accumulibacter” contain NulOs and how to enrich and/or isolate them. We extracted the EPS from granules enriched with “*Ca*. Accumulibcater” and used size-exclusion chromatography to separate them into different molecular weight fractions. This separation resulted in two high molecular weight (> 5,500 kDa) fractions dominated by polysaccharides, with a NulO content up to 4 times higher than the extracted EPS. This suggests that NulOs in “*Ca*. Accumulibacter” are likely located in high molecular weight polysaccharides. Additionally, it was seen that the extracted EPS and the NulO-rich fractions can bind and neutralize histones. This suggest that they can serve as source for sepsis treatment drugs, although further purification needs to be evaluated.

**Graphical abstract:** 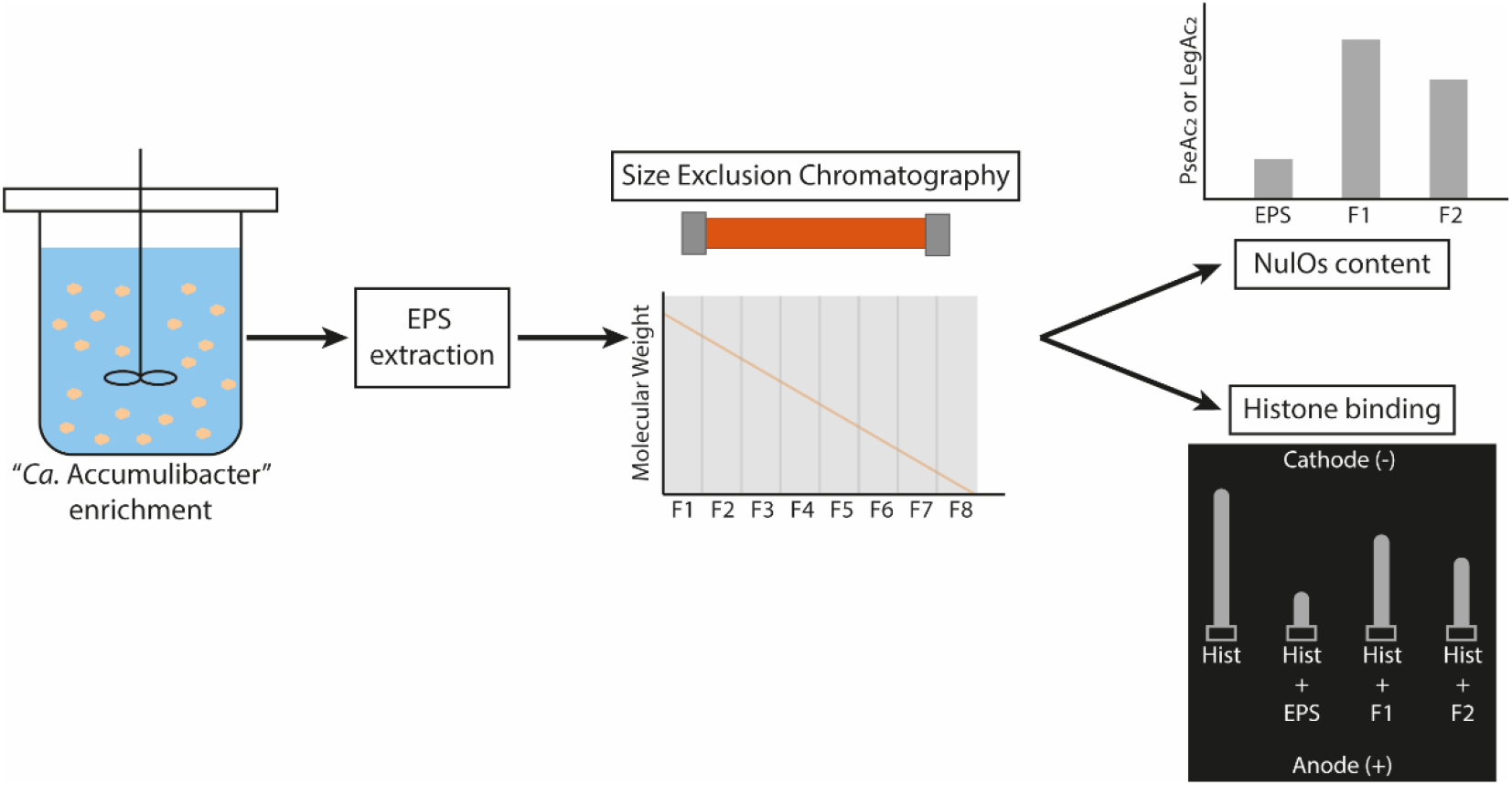

**Highlights:**

- NulOs in “Ca. Accumulibacter” are likely located in high molecular weight polysaccharides.
- Size exclusion chromatography allows to obtain high molecular weight polysaccharide-rich fractions enriched with NulOs.
- EPS and the NulOs-rich fractions can serve as source for sepsis treatment drugs.

## Introduction

Nonulosonic acids (NulOs) are a family of α-keto-acid carbohydrates with a nine-carbon backbone, with a wide variety of chemical forms. The different NulOs are normally found as terminal residues of extracellular glycoconjugates, acting as recognition molecules. The most studied representatives are derivatives of neuraminic (Neu) and ketodeoxynonulosonic (Kdn) acids (also known as sialic acids), specially N-acetyl-neuraminic acid (Neu5Ac), due to their importance in human physiology (Chen and Varki, 2010). However, there is a subgroup of NulOs that are unique to bacterial species. Examples of these are the derivatives of pseudaminic (Pse) or legionaminic (Leg) acids, which are often referred as “bacterial sialic acids” (Knirel et al., 2003). Bacteria have been reported to use these NulOs to decorate their surface polymers, such as capsular polysaccharides, lipopolysaccharides, flagella or S-layer glycocoproteins. Bacteria can also polymerize NulOs in these structures, forming polysialic acid (polySia) chains, with different degrees of polymerization (Haines-Menges et al., 2015). Bacterial sialic acids have been mainly studied for their role in pathogenesis. In pathogenic bacteria, these molecules serve as virulence factor and as mechanism of evading the host’s immune response by molecular mimicking, due to the structural similarities with human sialic acids (Varki et al., 2017). Bacterial sialic acids have also been suggested to play important roles in bacterial motility and biofilm formation (Goon et al., 2003; Jurcisek et al., 2005). However, further research is needed to fully understand the exact role of Pse and Leg and their derivatives.

An important reason for the lack of advances in this field is the lack of chemical access to NulOs in general, and Pse and Leg and their derivatives in particular. Neu5Ac has been traditionally synthesized chemically or extracted from natural sources. Engineered bacteria (*i*.*e*., *Escherichia coli*) have also been explored for its production. However, the complex structure of Pse and Leg makes their synthesis, production and commercialization difficult (Flack et al., 2020). The biosynthetic pathway of these carbohydrates is complex and requires several steps (Tomek et al., 2017). This makes enzymatic methods for the production of these compounds and derivatives too complex and their production would be very costly (Chidwick et al., 2021). Although, there have been advances in the chemical synthesis of Pse and Leg, the production yields are still low for a proper commercial production (Carter and Kiefel, 2018). Moreover, the dependency of organic solvents for their chemical synthesis is a concern for the sustainability of the production. Microbial biosynthesis of Pse and Leg has been mainly studied in pathogenic bacteria, which complicates the use of these organisms as production method. Therefore, new sustainable and efficient ways of production need to be explored.

A genome level study revealed that the biosynthetic pathway for different NulOs is widespread among archaea and bacteria (Lewis et al., 2009). However, NulOs have been overlooked in non-pathogenic bacteria. Very recently, a mass spectrometry based survey revealed an unexpectedly wide distribution of NulOs among non-pathogenic environmental bacteria (Kleikamp et al., 2020). Pinel et al. (2020) described the presence of Kdn and bacterial sialic acids in biofilms forming in cooling towers. In wastewater environments, Boleij et al. (2020) detected NeuAc, Kdn and bacterial sialic acids in the extracellular polymeric substances (EPS) of anammox granular sludge. Sialic acids were also identified in aerobic granular sludge dominated with “*Candidatus* Accumulibacter” (de Graaff et al., 2019). Further research with a highly enriched culture of “*Ca*. Accumulibacter” revealed its potential to produce different type of NulOs as part of its EPS, primarily the bacterial sialic acids Pse and/or Leg, which have the same molecular weight (Tomás-Martínez et al., 2021). Although the role of these carbohydrates in non-pathogenic environmental bacteria is still unknown, these findings point towards a new potential sustainable source of bacterial sialic acids.

“*Ca*. Accumulibacter” is the most abundant and well-studied polyphosphate accumulating organism (PAO) in wastewater treatment plants with biological phosphorus removal. Even though this microorganism has never been isolated, it has been successfully cultivated in laboratory bioreactors for decades (Smolders et al., 1994), reaching levels of enrichment of more than 95 % (Guedes da Silva et al., 2020). This successful enrichment has been achieved by employing ecological selection principles. “*Ca*. Accumulibacter” grows in the form of compact bioaggregates (granules) held together by the EPS (Barr et al., 2016; Weissbrodt et al., 2013). If bacterial sialic acids can be produced by a natural enrichment of “*Ca*. Accumulibacter” in a mixed-culture bioreactor, the aforementioned problem of involving pathogenic bacteria in the production process and the high cost of employing pure cultures will be avoided. This will be beneficial for the large scale industrial production of NulOs.

In order to employ a “*Ca*. Accumulibacter” enrichment to produce bacterial sialic acids, it is necessary to determine which fractions of the EPS of “*Ca*. Accumulibacter” contain NulOs and how to enrich and/or isolate them. In addition, NulOs can be polymerized into polysialic acid chains, conferring a high negative charge density. The polyanionic characteristics of these polymers allows their application in binding and neutralizing positively charged compounds, such as against histone-mediated cytotoxicity. Positively charged histones act as antimicrobial peptides to combat against pathogens. However, they are also toxic for host cells and excessive extracellular histones are associated with the development of sepsis or other diseases (Xu et al., 2009). Negatively charged polysialic acid inactivate the cytotoxic characteristics of histones (Galuska et al., 2017; Zlatina et al., 2017).

The aim of the present research was to determine in which EPS component of “*Ca*. Accumulibacter” NulOs are by fractionation and to evaluate the potential application of the NulOs-rich fractions against sepsis. EPS were extracted and characterized from granules from a lab enrichment of “*Ca*. Accumulibacter”. Extracted EPS was separated into different molecular weight fractions using size-exclusion chromatography. The anionic characteristic of the fractions was evaluated and the NulOs content was measured. Finally, the potential application against sepsis was evaluated performing a histone-binding assay.

## Materials and methods

### Reactor operation and characterization

The PAO enrichment was obtained in a 2 L (1.5 L working volume) sequencing batch reactor (SBR), following conditions similar to the one described by Guedes da Silva *et al*. (2020) with some adaptations. The reactor was inoculated using activated sludge from a municipal wastewater treatment plant (Harnaschpolder, The Netherlands). Each SBR cycle lasted 6 hours, consisting of 20 minutes of settling, 15 minutes of effluent removal, 5 minutes of N_2_ sparging, 5 minutes of feeding, 135 minutes of anaerobic phase and 180 minutes of aerobic phase. The hydraulic retention time (HRT) was 12 hours (removal of 750 mL of broth per cycle). The average solids retention time (SRT) was controlled to 8 days by the removal of effluent at the end of the mixed aerobic phase. The pH was controlled at 7.0 ± 0.1 by dosing 0.2 M HCl or 0.2 M NaOH. The temperature was maintained at 20 ± 1 °C. The reactor was fed with two separate media: a concentrated COD medium (400 mg COD/L) of 68:32 g_COD_/g_COD_ acetate:propionate (5.53 g/L NaAc·3H_2_O, 1.20 g/L NaPr, 0.04 g/L yeast extract) and a concentrated mineral medium (1.53 g/L NH_4_Cl, 1.59 g/L MgSO_4_·7H_2_O, 0.40 g/L CaCl_2_·2H_2_O, 0.48 KCl, 0.04 g/L N-allylthiourea (ATU), 2.22 g/L NaH_2_PO_4_·H_2_O, 6 mL/L of trace element solution prepared following Smolders et al. (1994). In each cycle, 50 mL of each medium was added to the reactor, together with 400 mL of demineralized water. The final feed contained 400 mg COD/L of acetate. Electrical conductivity in the bulk liquid was used to follow phosphate release and uptake patterns and to verify the steady performance of the reactor (Weissbrodt et al., 2014). Extracellular concentrations of phosphate and ammonium were measured with a Gallery Discrete Analyzer (Thermo Fisher Scientific, Waltham, MA). Acetate was measured by high performance liquid chromatography (HPLC) with an Aminex HPX-87H column (Bio-Rad, Hercules, CA), coupled to RI and UV detectors (Waters, Milford, MA), using 0.01 M phosphoric acid as eluent supplied at a flowrate of 0.6 mL/min.

### Microbial community analysis

The microbial community was characterized by 16S rRNA gene amplicon sequencing. DNA was extracted from the granules using the DNeasy UltraClean Microbial kit (Qiagen, Venlo, The Netherlands), using the manufacturer’s protocol. The extracted DNA was quantified using a Qubit 4 (Thermo Fisher Scientific, Waltham, MA). Samples were sent to Novogene Ltd. (Hong Kong, China) for amplicon sequencing of the V3-4 hypervariable region of the 16S rRNA gene (position 341-806) on a MiSeq desktop sequencing platform (Illumina, San Diego, CA) operated under paired-end mode. The raw sequencing reads were processed by Novogene Ltd. (Hong Kong, China) and quality filtered using the QIIME software (Caporaso et al., 2010). Chimeric sequences were removed using UCHIME (Edgar et al., 2011) and sequences with ≥97% identity were assigned to the same operational taxonomic units (OTUs) using UPARSE (Edgar, 2013). Each OTU was taxonomically annotated using the Mothur software against the SSU rRNA database of the SILVA Database (Quast et al., 2013).

### EPS extraction

Biomass samples collected at the end of the aerobic phase were freeze-dried prior to EPS extraction. EPS were extracted in alkaline conditions at high temperature, using a method adapted from Felz et al. (2016). Freeze-dried biomass were stirred in of 0.1 M NaOH (1 % w/v of volatile solids) at 80 °C for 30 min. Extraction mixtures were centrifuged at 4000xg at 4 °C for 20 min. Supernatants were collected and dialyzed overnight in dialysis tubing with a molecular cut-off of 3.5 kDa, frozen at −80 °C and freeze-dried. The freeze-dried extracted EPS samples were stored for further analysis.

### Size-exclusion chromatography fractionation

Freeze-dried EPS was solubilized in NaOH 0.01 M to a concentration of 10 mg/mL. Size-exclusion chromatography (SEC) was performed using a Hiload 16/600 Superose 6 prepacked column (Cytiva Lifesciences, Marlborough, MA) fitted on a system containing GX-271 dispenser/dilutor, a 307 pump and a 112 UV (280 nm) detector (Gilson, Middleton, WI). Fractions of molecular weights (MW) were determined after calibration with a HMW gel filtration calibration kit (44-669 kDa) (Cytiva Lifesciences, Marlborough, MA) and Blue Dextran (2,000 kDa). Molecular weights higher than the standards were calculated by linear extrapolation of the calibration line. A total of 15 mL of dissolved EPS was injected in the column with a flow rate of 1 mL/min. The running buffer consisted of 0.15 M NaCl and 0.05 M glycine-NaOH at pH 10. Seven different fractions were collected with MW ranges as shown in Table 1. The different fractions were dialyzed overnight in dialysis tubing with a molecular cut-off of 3.5 kDa, frozen at −80 °C and freeze-dried. The freeze-dried fractions were stored for further analysis.

**Table 1.** Weight distribution of the fractions obtained from SEC of the extracted EPS after dialysis and lyophilization. The non-soluble fraction represents the remaining solids after solubilization of the extracted EPS prior to the fractionation.

| Fraction Name | MW range (kDa) | Weight percentage of EPS (%) |
| --- | --- | --- |
| F1 | >15,000 | 7.7 |
| F2 | 5,500-15,000 | 8.4 |
| F3 | 738-5,500 | 9.8 |
| F4 | 100-738 | 12.1 |
| F5 | 12-100 | 8.9 |
| F6 | 5-12 | 34.7 |
| F7 | 3-5 | 5.7 |
| Non-soluble fraction |  | 12.7 |

### EPS and fractions characterization

#### Protein and carbohydrate content

Protein content was estimated using the bicinchoninic acid (BCA) assay (Smith et al., 1985) with bovine serum albumin (BSA) as standard. Carbohydrate content was determined using the phenol-sulfuric acid assay (Dubois et al., 1956) with glucose as standard. Both methods were used as described by Felz et al. (2019).

#### Fourier-transformed infra-red (FT-IR) spectroscopy

The FT-IR spectra of the different fractions was recorded on a FT-IR spectrometer (Perkin Elmer, Shelton, CT) at room temperature, with a wavenumber range from 550 to 4000 cm^-1^. Resolution of 1 cm^-1^ and accumulation of 8 scans were applied to each sample.

#### Nonulosonic acid analysis

NulOs were analyzed by high resolution mass spectrometry according to Kleikamp et al. (2020). Freeze-dried biomass were hydrolyzed in of diluted (2 M) acetic acid during 2 hours at 80 °C. After centrifugation, samples were dried using a Speedbac concentrator and labelled using DMB (1,2-diamino-4,5-methylene dioxybenzene dihydrochloride) during 2.5 hours at 50 °C. Labelled NulOs were analyzed by reverse phase chromatography Orbitrap mass spectrometry (QE plus quadrupole Orbitrap, Thermo Fisher Scientific, Waltham, MA). NulOs were identified according to their mass. To estimate the relative amount of NulOs in the samples, the peak area of a standard of Kdn was used as reference.

#### SDS-PAGE analysis and staining with Alcian Blue

SDS-PAGE was performed using NuPage® Novex 4-12 % Bis-Tris gels (Invitrogen, Waltham, MA) as described by Boleij et al. (2018). After dissolving in NaOH 0.1 M, each fraction was prepared in NuPAGE LDS-buffer and DTT (dithiothreitol) was added to a final concentration of 10 mM. Samples were incubated at 70 °C for 10 min for protein denaturation. A volume of 10 µL of sample was loaded per well. The Spectra Multicolor Broad Range Protein Ladder (Thermo Fisher Scientific, Waltham, MA) was used as MW marker. Gel electrophoresis was performed at 200 V for 35 min. After electrophoresis, the gel was stained with Alcian Blue at pH 2.5 for the visualization of carboxylate groups (R-COO^-^). The gel was extensively washed in solution I (25 % (v/v) ethanol and 10 % (v/v) acetic acid) for 2.5 hours, refreshing the solution 4 times. After washing, the gel was stained in 0.125 % (v/v) Alcian Blue in solution I for 30 min and washed in solution I overnight.

### Histone binding and agarose gel electrophoresis

Interaction of EPS and the obtained fractions with histones was tested using a method adapted from Zlatina et al. (2017). A mass of 5 µg of histones (H1, H2A, or H2B) was incubated with different amounts of EPS (in a ratio of 1:1, 1:2 or 1:3 histone:EPS), fractions (in a ratio of 1:2, 1:3 or 1:4 histone:fraction) or free Neu5Ac (in a ratio 1:3 histone:Neu5Ac) in 50 mM Tris for 1 hour at 30 °C and 300 rpm. Subsequently, 1 µL glycerol was added to each sample and samples were loaded on a 0.8 % agarose gel in 500 mM Tris/HCl, 90 mM boric acid, pH 8.5. The electrophoresis was performed at 80 V for 90 min with a running buffer (90 mM Tris/HCl, 90 mM boric acid, pH 8.5). The agarose gel was stained with Coomassie Blue for 1 hour and washed in demineralized water overnight.

## Results

### EPS extraction

For this study, the EPS of a lab-scale enrichment of “*Ca*. Accumulibacter” performing phosphate removal were extracted. The reactor performance and microbial community composition was similar as in earlier reports (Guedes da Silva et al., 2020; Oehmen et al., 2005), showing high PAO activity and enrichment of “*Ca*. Accumulibacter” (Figure S1). The EPS extraction yield was 58.3±14.7 % w/w of volatile solids. The protein and carbohydrate content of the extracted EPS accounted for 60.7±6.8 and 19.0±4.3 % w/w of volatile solids of EPS, respectively. Due to the limitation of the total carbohydrate assay, NulOs are not detected by this method (de Graaff et al., 2019). Thus, the amount of NulOs was not included in the carbohydrate content of EPS.

### EPS fractionation and characterization

The extracted pool of EPS was solubilized and fractionated in different molecular weight (MW) ranges using size-exclusion chromatography (SEC). Extracted EPS was separated in seven different fractions with apparent molecular weights ranging from 3 to more than 15,000 kDa. Notably, part of the EPS could not be solubilized and was not injected for the fractionation (non-soluble fraction). Table 1 shows the contribution of each fraction to the overall EPS. Most of the fractions (F1-F5) contributed similarly, with weight percentages ranging from 7.7 to 12.1 %, with the exception of the smaller fractions. F6 (12-100 kDa) showed the highest contribution, corresponding to 34.7 % of the total extracted EPS. On the other hand, F7 represented the lowest amount and was excluded from the subsequent analyses due to insufficient sample.

It is worth pointing out that, the fractionation range of the Hiload 16/600 Superose 6 column used in this research is between 5 kDa to 5,000 kDa, and the elution limitation is 40,000 kDa. Although the molecular weight of fraction F1 and F2 are out of the fractionation range of the column, as it is still within the elution limitation, they were collected and analyzed. Their molecular weight range was calculated by extrapolating the calibration curve. Moreover, one should be aware that all the molecular weight specification of the column corresponds to “globular proteins”. As the EPS may not be globular proteins, the molecular weight measured by SEC can only be considered as a relative value.

For each of the fractions, the contents of total proteins and carbohydrates were estimated using colorimetric methods and BSA and glucose as standards, respectively. Figure 1 shows the carbohydrate to protein ratio for the extracted EPS and each of the fractions obtained from SEC. Although extracted EPS was dominated by proteins, the high MW fractions (F1 and F2) were dominated by carbohydrates (PS/PN ratio > 1). The decrease of the MW in the fractions was accompanied with an increase of protein content relative to the carbohydrate content. The smallest MW fractions (F4-F6) were dominated by proteins (PS/PN ratio < 1). Interestingly, F5 and F6 were mainly composed of proteins and the carbohydrate fraction was negligible (PS/PN ratio of 0.01). Thus, SEC allowed the separation of the extracted EPS in high MW carbohydrates dominated fractions.

**Figure 1.**
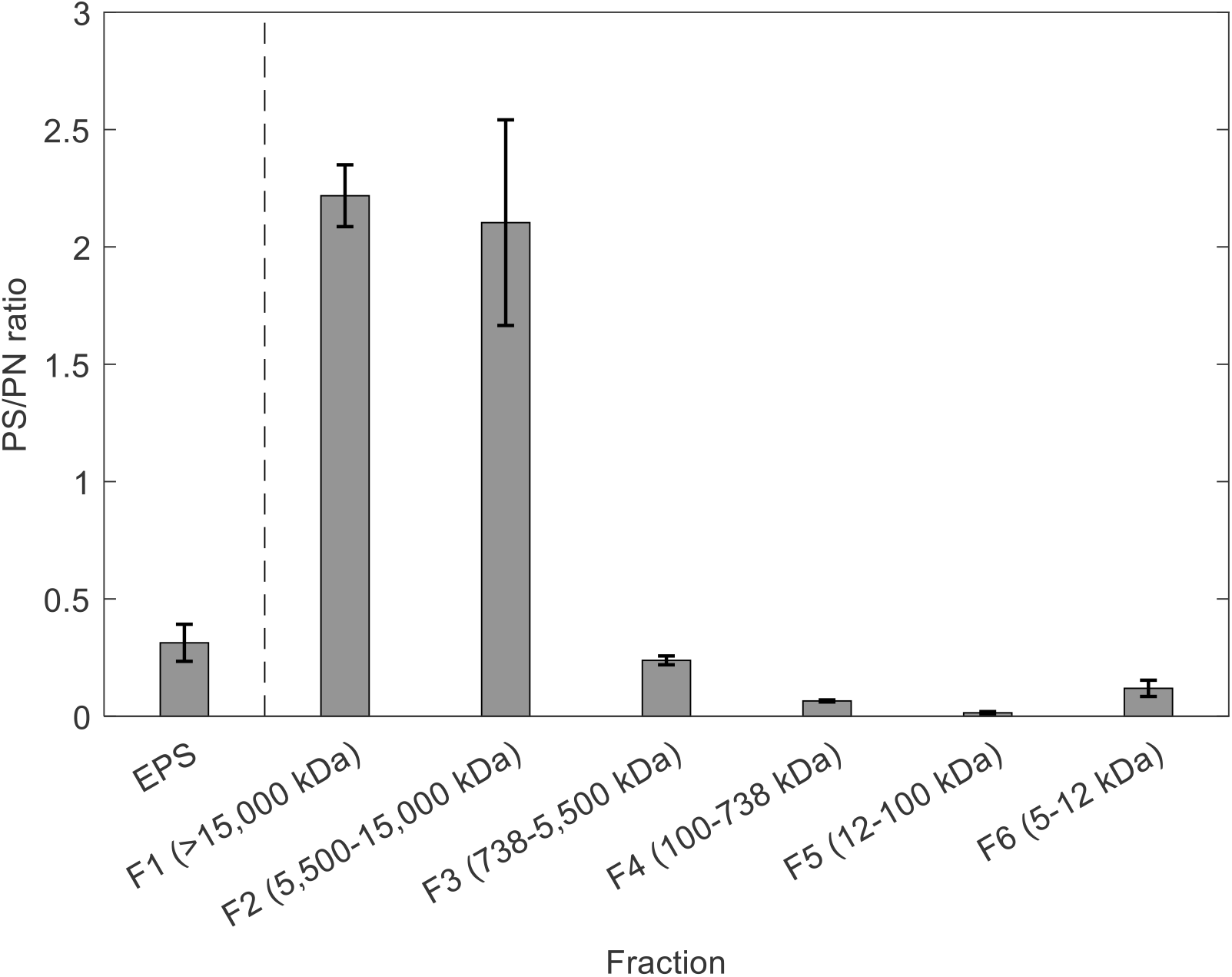
Carbohydrate to protein ratio (PS/PN) of the different MW fractions obtained by SEC. Carbohydrate and protein content are expressed as glucose and BSA equivalents.

In order to get a better evaluation of the differences between the obtained fractions, FT-IR spectroscopy was used to analyze their composition. Figure 2 shows the individual FT-IR spectrum of each of the fractions. These results confirmed the decrease of carbohydrate content and increase of protein content as the MW decreases. F1 shows a high peak at ∼1030 cm^-1^, corresponding to the C-O stretching of carbohydrates. This peak decreases in F2 and becomes negligible in the rest of the fractions. The opposite tendency occurs with the peaks at ∼1530 and ∼1640 cm^-1^, corresponding to the N-O stretching and N-H bending of proteins, respectively. This peak becomes dominant in the fractions with the lowest PS/PN ratio (F4 and F5). Additionally, F1 shows a peak at ∼1730 cm^-1^, which is associated to the α-keto aldonic acid structure of NulOs (de Graaff et al., 2019). This peak appears subtly in the spectrum of F2 and it is absent in the rest of the fractions. These results suggest the presence of NulOs in the high MW fractions (F1 and F2).

**Figure 2.**
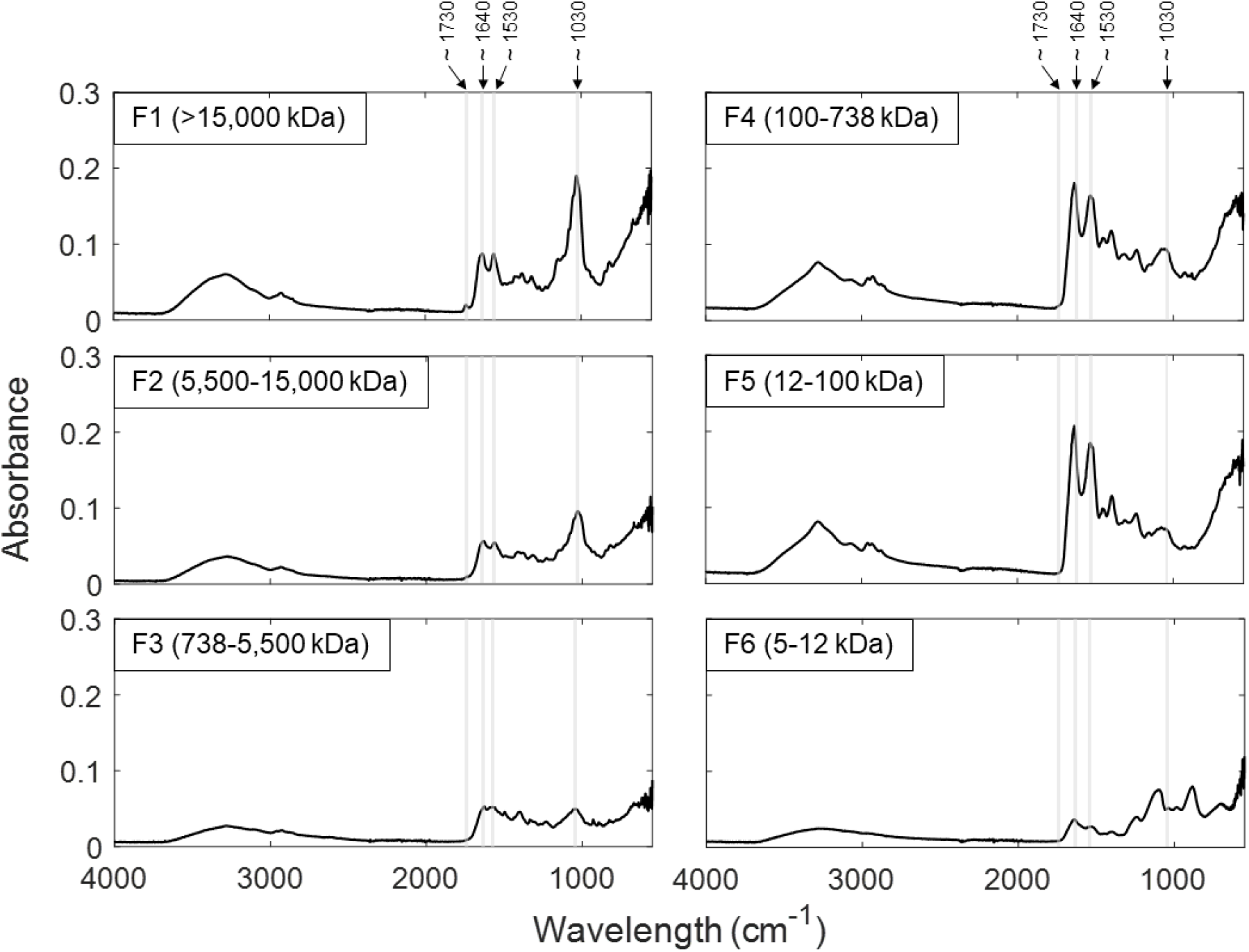
FT-IR spectra of the different MW fractions obtained by SEC. Gray areas highlight peaks corresponding to carbohydrates (∼ 1030 cm_-1_), proteins (∼ 1530 and ∼1640 cm_-1_) and NulOs (∼ 1730 cm_-1_).

To confirm the presence and type of NulOs in the EPS and the fractions obtained by SEC, the different samples were analyzed using mass spectrometry. It revealed the presence of double acetylated Pse or Leg (PseAc_2_ or LegAc_2_), which cannot be distinguished as they have the same molecular mass. This NulO was detected in the extracted EPS and in the high MW fractions (F1, F2 and F3). The rest of the fractions showed negligible amount of NulO. In order to estimate the amount of NulO in each sample, the area of PseAc_2_ or LegAc_2_ was compared to a reference amount of standard Kdn. Although this cannot be used as absolute quantification, it can give a relative estimate of the NulO content of each sample. The estimated content of PseAc_2_ or LegAc_2_ of each sample is given in Figure 3. The fractions F1 and F2 showed a higher content of PseAc_2_ or LegAc_2_ than the original EPS (4 and 3 times higher, respectively). The fractionation with SEC allowed to obtain fractions highly enriched with PseAc_2_ or LegAc_2,_. Those fractions are also carbohydrate-rich and with a MW >5,500 kDa.

**Figure 3.**
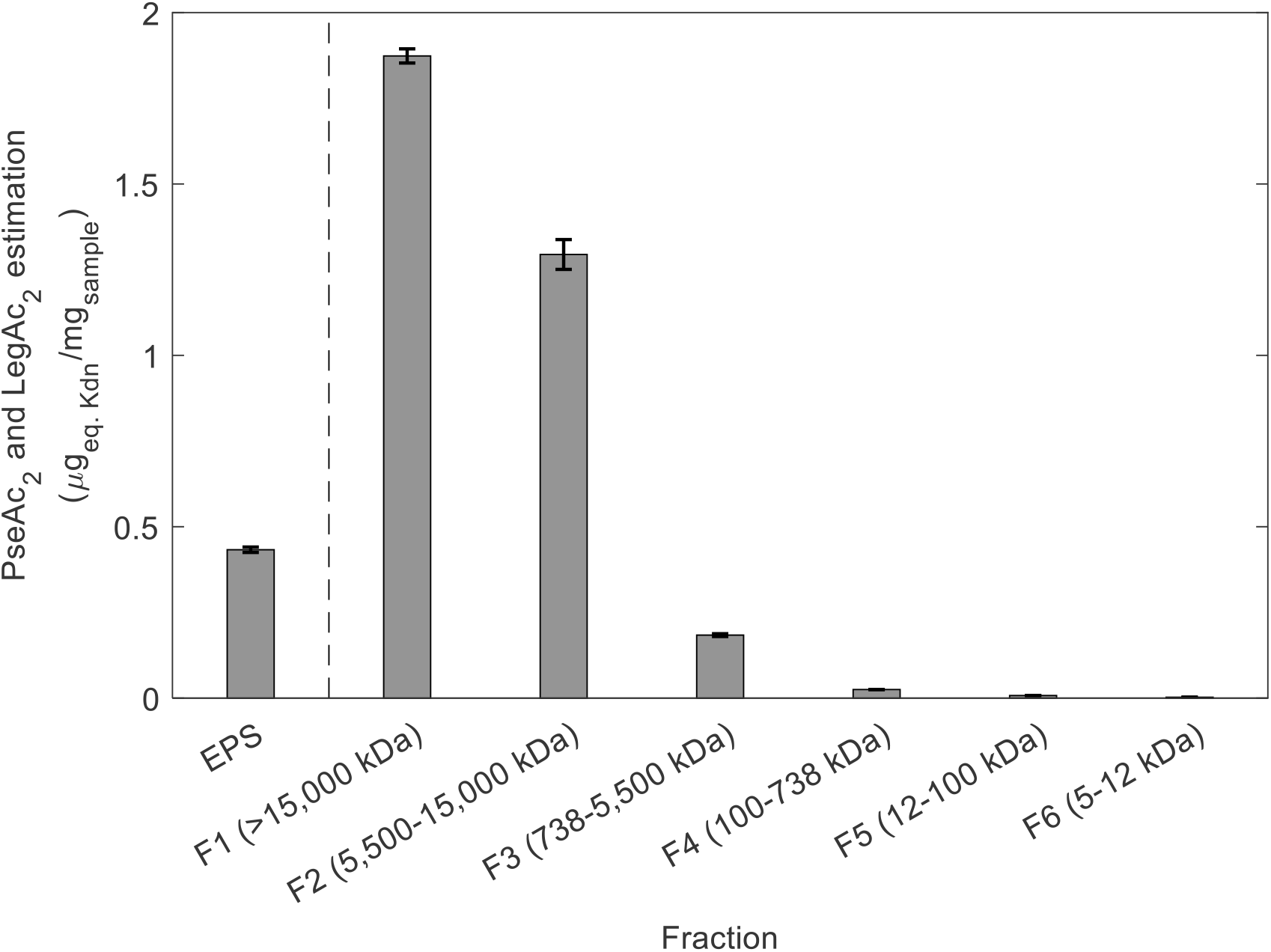
Relative quantification of NulOs in the extracted EPS and in the different MW fractions obtained by SEC. The detected NulO is LegAc_2_ or PseAc_2_, which could not be distinguished as the have the same molecular weight. The amount of NulOs was estimated based on the relative area of a spiked standard of Kdn.

The carboxylic group of NulOs can confer a negative charge to the polymer, which binds with Alcian Blue. In order to confirm and visualize the strongly acidic carboxylic groups in the extracted EPS and the separated fractions, samples were loaded in a SDS-PAGE gel. After the separation, the gel was stained using Alcian Blue at pH 2.5, which is specific for acidic glyconjugates (Figure 4). The extracted EPS and fractions F1 and F2 were heavily stained at the position corresponding to high molecular weight, implying the presence of acidic glycoconjugates. In comparison, F3 was slightly stained and F4-F6 were not stained at all. This confirmed the presence of strong acidic groups in the high MW fractions, which is in line with the high amount of PseAc_2_ or LegAc_2_ in these fractions.

**Figure 4.**
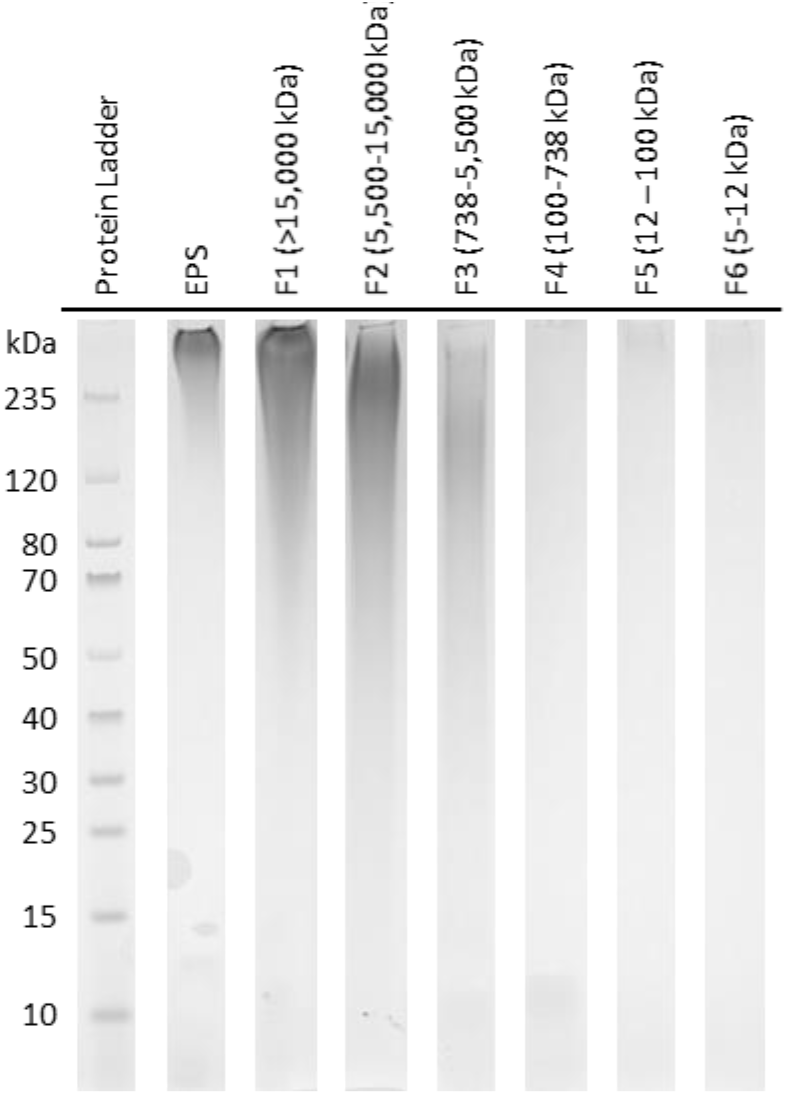
SDS-PAGE gel of the extracted EPS and the different MW fractions obtained by SEC. The gel was stained with Alcian Blue at pH 2.5 for acidic carbohydrates.

### Histone binding assay

Negatively charged polymers, such as polysialic acids or heparin, can be used as treatment of sepsis due to their capacity to bind histones. To test the potential of extracted EPS and the NulO-rich fractions (F1 and F2) for application in the treatment of sepsis, a histone binding assay was performed. Histones were incubated with the different samples and the migration characteristics was evaluated (Figure 5). Histones (*e*.*g*., H1, H2A and H2B) are positively charged and they migrate towards the cathode (negative pole). When they are incubated with negatively charged polymers, if the interaction result to the neutralization of the charge of histones, their migration towards the cathode will be reduced.

**Figure 5.**
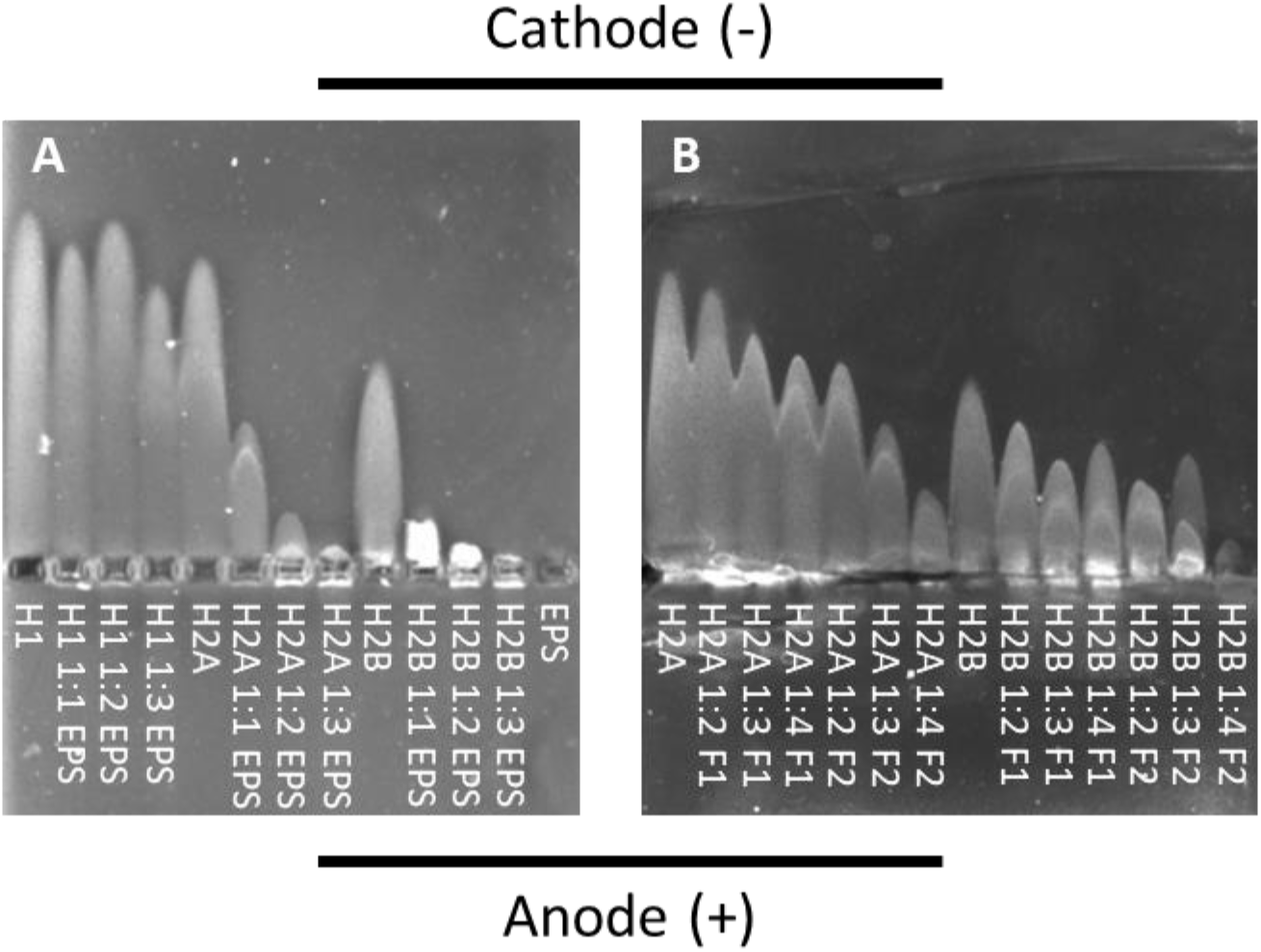
Histone binding assay. Histones were incubated with different amounts (as indicated by the mass ratio) of extracted EPS (A) or NulO-rich fractions (F1 (>15,000 kDa) and F2 (5,500-15,000 kDa)) (B). After electrophoresis, gels were stained with Coomassie Blue.

Firstly, the histone-binding capacity of EPS was tested with three different histones (H1, H2A and H2B) by dosing different amounts of EPS (Figure 5A). In the case of histone H1, the migration was only slightly reduced when a dosage ratio of 1:3 histone:EPS was used. When a lower dosage of EPS was used, no migration reduction was observed. In the case of histones H2A and H2B, a dosage ratio of 1:1 histone:EPS was already effective, as the migration was only half of the histone control. An increase of dosage ratio to 1:2 and 1:3, significantly decreased the migration characteristics of the histones.

As H2A and H2B are the most abundant histones causing sepsis (Zlatina et al., 2017), they were further tested with the NulO-rich fractions (F1 and F2) (Figure 5B). Both fractions reduced the migration of the histone H2B but only F2 reduced the migration of histone H2A. This indicated that they bind with the histones and neutralized their charge. Once the dosage ratio was increased from 1:2 to 1:4 histone:EPS, the neutralization effect increased as well. Generally, the neutralization effect of both NulO-rich fractions is stronger with H2B than with H2A. F2 had a higher reduction of the migration of both histones, when compared to F1, even though the NulO content was lower in F2. It was also noticed that the binding capacity of the extracted EPS was higher than the NulO-rich fractions, a dosage ratio of 1:3 histone:EPS already inhibited the migration of both histones. The exact interaction between EPS (including the fractions) and histones is unclear: probably the polymer conformation or the polymerization degree of NulOs play a role in addition to the charge, or there are other binding sites besides the NulO, for example sulfated glycosaminoglycans.

## Discussion

### An enrichment of “*Ca*. Accumulibacter” can be a potential production platform of Pse or Leg derivatives

The lack of chemical access to Pse and Leg and their derivatives hinders the study of these type of NulOs. Although their production has been attempted through engineered bacteria or chemical synthesis (Carter and Kiefel, 2018; Flack et al., 2020), it is still very costly and too complex. A sustainable alternative can be the use of mixed cultures, which require non-sterile conditions and reduces the cost substantially (Kleerebezem and van Loosdrecht, 2007). This mixed culture biotechnology has been employed for the production of different chemicals, such as, polyhydroxyalkanoates, organic acids or medium-chain fatty acids, among others (Dionisi and Silva, 2016; Serafim et al., 2008; Stamatopoulou et al., 2020).

Recently, it was demonstrated that “*Ca*. Accumulibacter” has the potential to produce different types of NulOs in its EPS (Tomás-Martínez et al., 2021). In the current work, we showed the production of PseAc_2_ or LegAc_2_ by “*Ca*. Accumulibacter” under our reactor conditions. This bacteria can be cultivated in lab reactors, reaching dominance levels of up to 95 % of the total community (Guedes da Silva et al., 2020). This type of reactors allows a good control of the conditions, ensuring the long term reproducibility of the production. Moreover, culture conditions can be easily adapted for specific requirements. For example, it could be determined under which conditions the NulOs production is optimized, and by manipulating the operational conditions, to reach the optimized production.

After NulOs production by “*Ca*. Accumulibacter”, a purification strategy needs to be implemented. Here, we managed to increase the NulO content 4 times by a simple purification. Firstly, EPS was extracted by alkaline conditions. The resulting extracted EPS was then fractionated based on molecular weight using size-exclusion chromatography (SEC). This resulted in a NulO enrichment in the fractions corresponding to the highest molecular weights (>5,500 kDa). Further purification has to be explored to obtain PseAc_2_ or LegAc_2_. Mild acetic acid hydrolysis has been used previously to release and purify Pse from polysaccharides (Lee et al., 2018). After the release, additional separation steps would be needed to obtain the final product.

### NulOs in “Ca. Accumulibacter” are likely located in high MW carbohydrates

The separation of the extracted EPS into different MW fractions revealed that PseAc_2_ and/or LegAc_2_ are present in high MW polymers (F1 and F2). These fractions, as oppose to the rest, are rich in carbohydrates. Typically, polysaccharides have a much higher MW than proteins. Gómez-Ordóñez et al. (2012) described polysaccharides from seaweed with MW higher than 2,400 kDa. Liu et al. (2016) showed the presence of polymers with MW higher than 1,000 kDa in the EPS of aerobic granules. However, most of the studies in bacterial EPS report a MW to a maximum of 670 kDa (Garnier et al., 2005; Simon et al., 2009). It is noted that, the fractionation range and elution limitation of the SEC column in those studies were much lower than the current research. Probably due to the separation limitation of the column, EPS with higher molecular weight was overlooked. On the other hand, glycosylation of proteins can significantly increase their apparent molecular weight in SEC separation. Human mucus is a complex polymeric mixture with protein biomolecules ranging from 6 kDa to 100 MDa. Specifically, mucins have a typical MW of 200 kDa to 100 MDa (Radicioni et al., 2016). Mucins are highly glycosylated proteins linked with sialic acids and represent 20-30 % by weight of the mucus. As the PS/PN ratio in fractions F1 and F2 is higher than 1, there is possibility that these two fractions are highly glycosylated proteins comparable to mucins, with similar MW range (5.5 MDa – 40 MDa), linked with bacterial sialic acids.

Pathogenic bacteria have been described to decorate some of their surface polymers with NulOs, such as capsular polysaccharides, lipopolysaccharides, flagella or S-layer glycocoproteins (Haines-Menges et al., 2015). The MW of these polymers has been reported to range from tens to hundreds kDa. The capsular polysaccharide of *Streptococcus pneumonia* ranged from 606 to 1,145 kDa (Bednar and Hennessey, 1993). For some strains of *E. coli*, the described MW was lower, ranging from 51.3 to 130.6 kDa (Restaino et al., 2019). Their results showed that lipopolysaccharides have a higher MW than capsular polysaccharide, judging from their elution time in SEC.

Although it was not determined exactly which type of polymer contains PseAc_2_ or LegAc_2_ in “*Ca*. Accumulibacter”, definitely the NulO-containing polymer is highly glycosylated with a high MW. This could be a glycoprotein similar to mucins, or lipopolysaccharides with a high carbohydrate content. Further purification and analysis could reveal the exact location of these NulOs.

### NulOs-rich EPS and fractions as potential source for sepsis treatment drugs

Negatively charged polysaccharides such as heparin or polysialic acids have a cytoprotective effect by neutralization of extracellular histones (Ulm et al., 2013; Wang et al., 2020b). We demonstrated the potential use of the extracted EPS from “*Ca*. Accumulibacter” for this application. The extracted EPS can bind histones H2A and H2B and neutralize them, as indicated by the decrease of migration distance in Figure 5. However, the complex composition of EPS will hinder their direct application in the medical field. Separation and purification techniques are needed to obtain compounds that can act as final sepsis treatment drugs.

In this study, the use of SEC for the separation of the extracted EPS allowed to obtain fractions rich in NulOs and dominated by polysaccharides (F1 and F2). For medical application, polysaccharides are preferred over proteins, as proteins show stability and immunogenicity problems (Wang et al., 2020a). Fractions F1 and F2 were tested for their capacity to neutralize histones. A slightly lower capacity than the extracted EPS was observed, which can be compensated by a higher dosage to achieve a similar effect as the extracted EPS. Although F1 had a higher NulOs content than F2, F2 had a higher histone neutralization effect. Additionally, the higher the dosage, the stronger the effect was. According to Zlatina et al. (2017), polysialic acids rather than single sialic acid monomer manifest the neutralization capacity to histones. Moreover, this capacity of polysialic acids depends on the degree of polymerization. This might explain the higher effect of F2, where PseAc_2_ or LegAc_2_ might have a higher degree of polymerization than in F1.

It was noticed that the extracted EPS displayed a stronger neutralizing effect than the NulOs-rich fractions. These differences could be caused by the presence of binding sites other than NulOs in the EPS. For instance, sulfated polysaccharides have been described in the EPS of aerobic and anaerobic granular sludge (de Bruin et al., 2022; Felz et al., 2020), which could potentially contribute to the histone binding capacity of EPS. Further research is needed to examine all the histone binding sites in the EPS and the fractions, in order to fully understand the neutralization mechanisms.

## Conclusion

In this study we showed that enrichments of “*Ca*. Accumulibacter” can be a potential sustainable alternative for the production of bacterial NulOs (*e*.*g*., PseAc_2_ or LegAc_2_). Size-exclusion chromatography equipped with high molecular weight separation column can be used as initial purification step to obtain NulOs-rich fractions. This separation obtained high molecular weight fractions (> 5,500 kDa) dominated by polysaccharides, where the NulO content was increased up to 4 times, compared with the extracted EPS. Additionally, the capacity of EPS and these fractions to bind histones suggest that they can serve as source for sepsis treatment drugs, although further purification needs to be evaluated.

