## Supplementary figures and images for "Enrichment and application of bacterial sialic acids containing polymers from the extracellular polymeric substances of “*Candidatus* Accumulibacter”"

### Figure S1

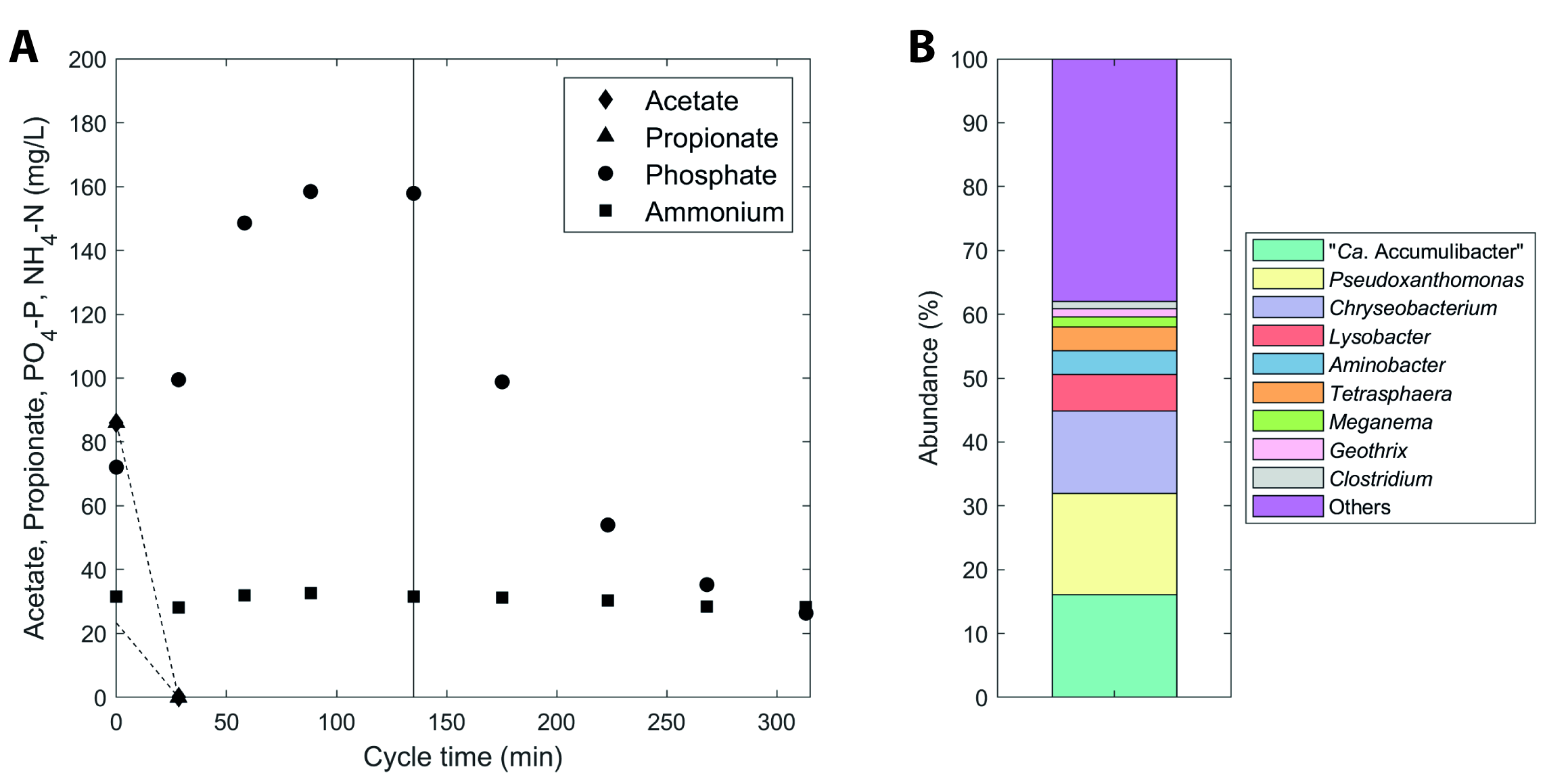
